# “Light-box” accelerated growth of poinsettias: LED-only Illumination

**DOI:** 10.1101/264168

**Authors:** Charitha Weerasuriya, Stewart Detez, Soon Hock Ng, Andrew Hughes, Michael Callaway, Iain Harrison, Tomas Katkus, Saulius Juodkazis

## Abstract

For the current commercialized agricultural industry which requires a reduced product lead time to customer and supply all year round, an artificial light emitting diode (LED)-based illumination has high potential due to high efficiency of electrical-to-light conversion. The main advantage of the deployed Red Green Blue Amber LED lighting system is colour mixing capability, which means the ability to generate all the colours in the spectrum by using three or four primary colour LEDs. The accelerated plant growth was carried out in a “light-box” which was made to generate an artificial day/night cycle and had a capability to tune the colour mixing ratio along the colour temperature curve of the chromaticity diagram. The control group of plants form the same initial batch was grown on the same shelf in a greenhouse at the same conditions with addition of artificial illumination by incandescent lamps for few hours. More dense and better quality (hence, commercial value) plants were grown in the light-box. Costs and efficiency projections of LED lamps for horticultural applications is discussed together with required capital investment. The total cost of the “light-box” including LED lamps and electronics was 850 AUD.

## 1. INTRODUCTION

Light-emitting diodes (LEDs) are the most efficient lighting devices and used in many luminaire applications. Luminaires with LED chips have flexibility to control its light output as per the application requirements, including colour temperature, colour spectrum, and saturation. Combination of four primary colours (red, green, blue, amber) or RGBA per the CIE 1931 colour space chromaticity diagram can most reliably simulate the natural colours we see in nature.^1, 2^ Due to low power consumption and efficient light controlling methods, LED chips generate less heat compared to conventional high-pressure sodium lamps used in agriculture.^3^ Moreover, the lifetime of LED lamps of ~ 10 years makes further impact for competitiveness of LED lighting applications. This introduces new engineering design capabilities in placing lamps in proximity to the plants and allows the use vertical organisation of plants on shelves.

The main objective of the current LED lighting for plant growth applications is to investigate behaviour of the plant growth under artificial lighting. One example application where a circadian rhythm for a plant should be controlled is an accelerated industrial growth of Christmas poinsettia (*euphorbia pulcherrima*) for blossoming (colour change). In Australia (southern hemisphere), poinsettia shoots are planted in June for blossoming half-a-year later in December. This corresponds to an unnatural winter-to-summer cycle change for the Mexican plant, originally from the northern hemisphere.

Here we report a study of poinsettia growth using LED artificial lighting. Full control over natural lighting (intensity and colour) by using LED lamps in a fully controlled greenhouse environment was achieved and an accelerated growth cycle of poinsettias has been demonstrated. The test patch was ~ 1 m^2^ and was fully separated from the outside greenhouse lights. One of the expected outcome of the project is to develop a light controlling system for accelerating plant growth.^4, 5^ The capital cost investment into the LED lighting system and the “light-box” for plant growth tests in a greenhouse is presented to facilitate estimation of technology readiness and competitiveness.

In particular for the tested plant poinsettia, *Euphorbia pulcherrima*, replication of shorter days triggering the colour change in the bracts (modified leaf stems) was tested under LED artificial illumination and altered day-night cycle.^6^ How specific spectrum and/or intensity effects for the bracts color change is currently not known.

### 2. SAMPLES AND PROCEDURES

### 2.1 Pointsettias

Pointsettia, *Euphorbia pulcherrima*, is a commercially grown plant, native to Mexico, well known for its red and green foliage, used as a floral display at Christmas time. The red colour isnt the flowers but coloured bracts located around the insignificant white/cream flowers.

The plants are purchased as plugs, small plants with well-established and independent root systems, from Ball Australia in August. The plugs are then planted into larger pots for growing on for sale pre-Christmas. Plants are sold in either singles or triples in each pot. The latter producing a larger fuller floral display.

The plants are grown in a controlled environment to produce a crop at its best ready for Christmas. Poinsettias are tropical plants and in Melbourne the crop begins its production process in winter. Therefore, they are grown within a controlled environment that reproduces the plants normal growing requirements, warmth and light. What triggers the plants bracts to change from a green to the bright red colour is the shorter periods of sunlight the plant experiences with the onset of winter. When the production crop of poinsettia plants have grown to a specified size, the plants are then exposed to shorter periods of sunlight, triggering the colour change in the bracts.^7^ The controlled growing space has a roof blind, that when closed, reduces the natural light the plants are exposed to, simulating shorting days and the onset of winter. The plants react, changing colour, producing the red blooms poinsettias are known for.

### 2.2 DIY: infrastructure, lights, and electronics

To design and develop a customized electronic product requires a lot of effort and funding. However, thanks to open source electronics modules, many kinds of electronic DIY projects are achievable within affordable budget. The light-box is one such DIY project attempted for this study and was designed by using open source off-the-shelf components.

The “light-box” of ~ 1 m^2^ was made out of light weight aluminium and covered with a reflective Al-foil. The coating aimed to homogenise the side-illumination of plants due to reflection from three LED lamps placed at the ceiling of the “light-box”. The “light-box” was 1200 mm (Length) × 800 mm (Width) × 900 mm (Height) made from Aluminium angle 40 mm × 40 mm × 1.4 mm and Aluminium flat bar 30 mm × 3 mm, pop rivets, reflective silver film (Ametelin, Silversark), insulation and ducting tape. An approximate cost of materials was 150 AUD and time to build 3 hours.

In the light box, the combined on-chip RGB LED strips have been used to produce all the colours in the visible spectrum. In addition, electronics included one 12 V, 2.5 A (up to 30 W power) constant voltage LED driver, Aurdino Uno module to control LED spectrum and communicate with the host, power metal oxide semiconductor field-effect transistor (MOS-FET) modules for switching and a light fixture of the standard fluorescent lamp.

The RGB LED strip contains 60 (Red), 60 (Green), 60 (Blue) chips per meter, and each individual chip is capable of generating 250 mcd, 880 mcd, 180 mcd red, green and blue luminous intensities (a luminous flux per unit solid angle 1 cd = 1 lm/srad), respectively, at nominal 20 mA current. Furthermore, wavelength of each colour is 640 nm Red, 525 nm Green, 470 nm Blue. In a single light fixture, three one-meter strips are used to increase luminous intensity.

Electronics and RGB-LEDs were bought online (Ebay). The light fixtures were adapted from fluorescent lamps (DIY Bunnings, Mercator Cannes lamp 32 AUD). Lamp diffuser was essential for color mixing by scattering (Fig. 1), however, caused some intensity loss. All electronics and power supply was placed in hermetic box and placed outside the “light-box”. The cost per LED lamp composed of three LED-ribbons and driver electronics was following: RGB LED-ribbon of 60-LEDs was 15 AUD, electronic drivers were 50 AUD.

**Figure 1.**
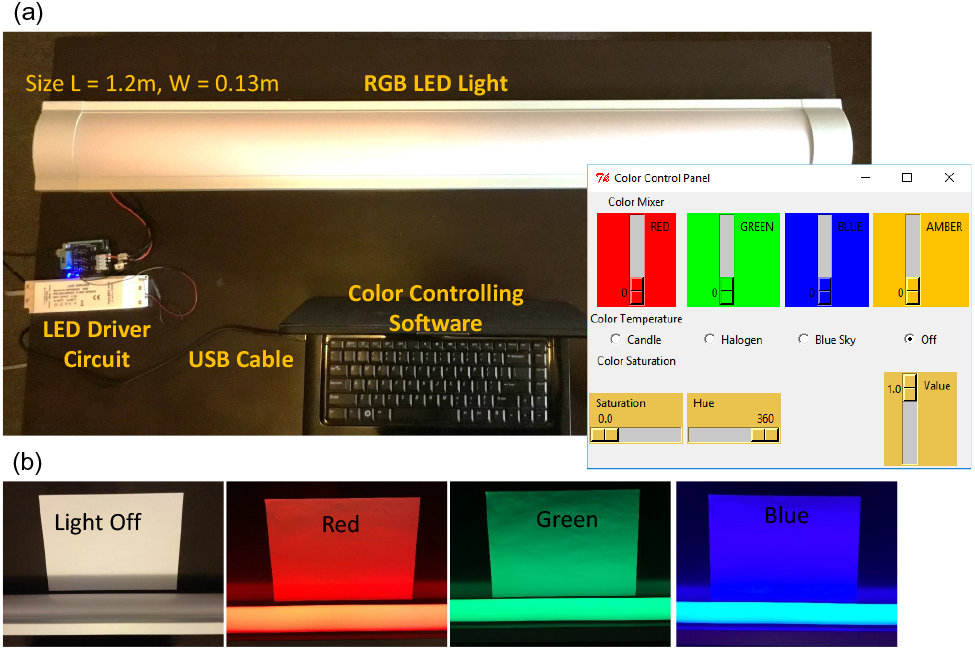
(a) Lamp with RGB LED-ribbons and controlling parts; separate single color-ribbons were used. Inset (right-side) shows the GUI for RGBA control of the LED-ribbons. (b) White paper sheet illuminated with different colors.

The experiment was controlled and data was collected using an in-house made program (Python coding) for color mixing, run from a portable computer (see the user interface in the inset of Fig. 1(a)). Once lighting conditions: color intensities, temporal exposure patterns were setup, the electronics ran independently.

For measurements of intensity and spectrum, a portable spectrometer Ocean Optics (OCEAN-FX-VIS-NIR) equipped with a 400-µm-core fiber (QP400-2-VIS-NIR) of numerical aperture *NA* = 0.22 *±* 0.02 was used.

Illuminance - a luminous flux incident on a surface - was measured with a hand-held luxmeter QM1587 (Digitech Ltd.) to compare different light flux in greenhouse and light-box; 1 lx = 1 lm/m^2^. e.g. a sunshine (not direct) amounts to 10-25 klx.

### 2.3 Color mixing algorithm

As explained above, the main idea is to control RGBA colours to generate visible spectrum. However, it is difficult to find RGBA LED strips or panels in the open electronic market. Therefore, RGB strips were selected at this initial state (Until A-amber ribbons are commercially available at the same price as the RGB). With RGBA colour, there are many parameters or configuration to set. At this moment, the focus is to monitor plant growth by manipulating day/night cycles. Day/night cycle can be simulated by controlling the correlated colour temperature of the chromaticity diagram.

Color rendering was realised following four-color RGBA color mixing,^1^ which allows the best coverage of color space and can achieve natural lighting, tailor more settle nuances of color dullness, intensity, hue and amplitude. Software to control LED lamp was written using Python coding and had RGBA functionality (Fig. 1(a)).

In this project, color rendering is based on varying spectral power between the green and red regions of CIE 1931 xy chromaticity diagram at the wavelength from 530 nm to 620 nm. The white light falls on AGB and RGB triangle.^1^ Hence, the white light can be generated with the mixture of spectral powers *S_AGB_* or *S_RGB_* or linear mixture of both by controlling red and amber color ratio:^1^

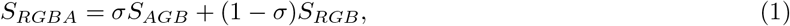

where *σ* is a weight parameter 0 <*σ*<1. Radiant flux is calculated by the following equations:

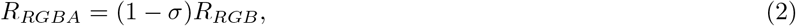

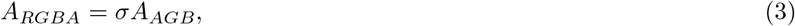

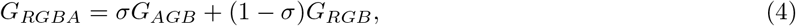

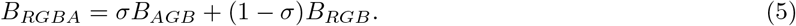

The rule above was used to set individual color intensity by the electrical current.

### 3. RESULTS AND DISCUSSION

LED lamps for experiments of plant growth under artificial illumination are commercially available (e.g., Hortiled, Ledigma Ltd.), however, at a prohibitively high cost (> 10k USD/m^2)^ for field experiments in a greenhouse. In Sec. 2 we showed price estimates which compound to 850 AUD for the ~ 1 m^2 “^light-box” (prices mid-2017).

### 3.1 Lighting conditions

Colour temperature can be realised using the RGB mixing. By manipulating colour temperature, it is possible to produce a wide range of white light output. For example, 1500 K to almost 10000 K depending on the LED chip quality. One of the controlling factors is to generate artificial day and night cycles to test plant response. The day and night feeling can be created by moving the colour mixing ratio along the colour temperature curve of the chromaticity diagram. For example, a colour temperature of 6000 K produces something similar to natural sky blue, cool white, while a colour temperature of 3000 K or less creates warm white. One of the main observation is to study how plant growth is affected by the day and night cycle.

For comparison of lighting conditions in the “light-box” and greenhouse, where commercial growth of poinsettias was carried out, we used a portable Ocean Optics spectrometer and a lux-meter. The optical fiber coupler has effective numerical aperture of *NA* = *n* sin *α* ≃ 0.22 and was collecting light form a half-angle of *α* ≃ 12.7° *n* = 1 is the refractive index of air. The corresponding solid angle was Ω = 2*π*(1 – cos *α*) ,..≃ 0.15 srad.

Figure 3(a) shows spectra measured in the “light-box” and in the greenhouse. Only RGB components are present in spectrum of the LED lamp compared with the spectrally broader spectrum from natural sun light inside the greenhouse. This makes a darker appearance of lighting conditions inside “light-box” (Fig. 2). The difference was quantified by calculating the area below the luminance curves (in Fig. 3(a)). Only ~ 25% of luminance was inside the light-box. Considering natural light cycle in the greenhouse and 24/7 illumination in the light-box, the difference in light dose reaching plant was more than twice. However, the rate of bio-mass production for plants under natural greenhouse illumination was only slightly faster as compared with the light-box where the dose of light received by plants was less than half.

**Figure 2.**
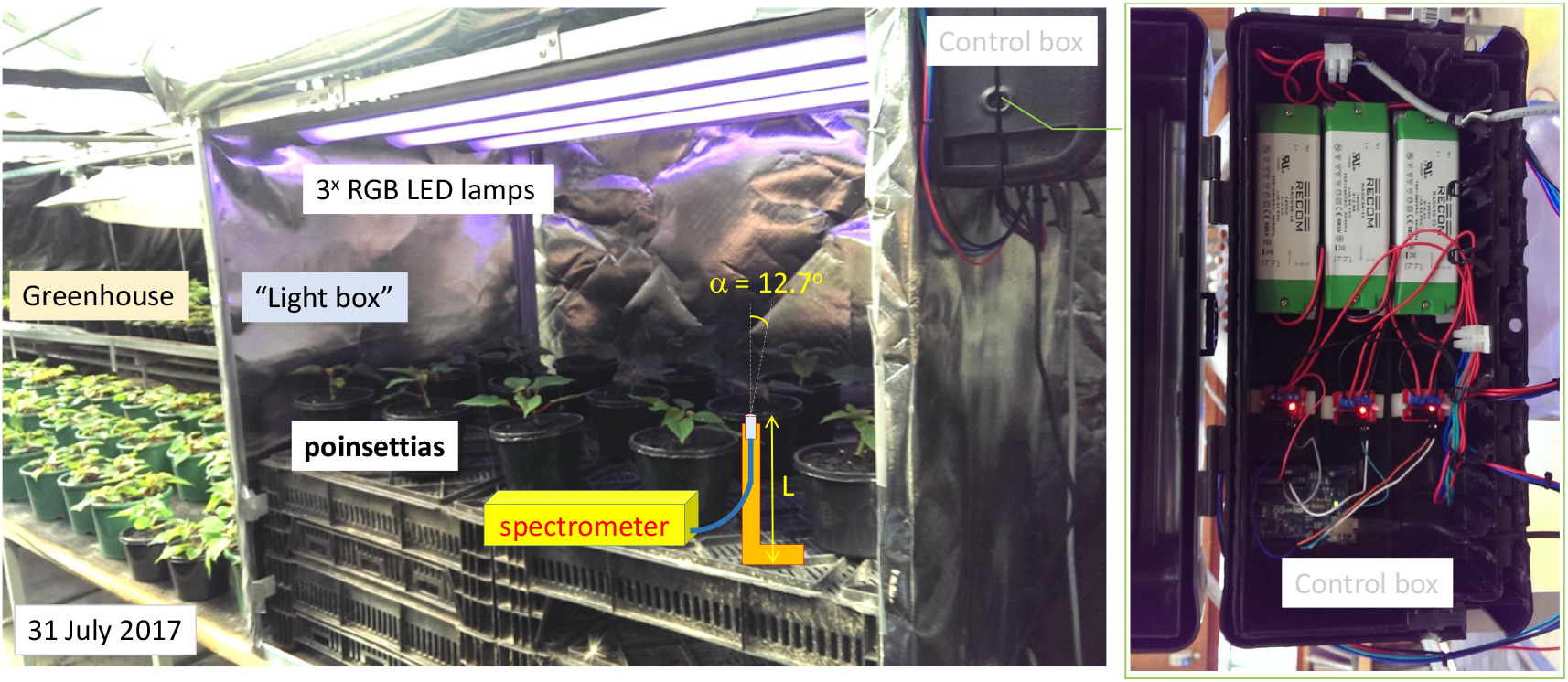
Start of experiment at Wantirna greenhouse using RGB LED lamp. Color spectrum was set in a “light-box”: Red (640 nm) 100%, Blue (470 nm) 100% and Green (525 nm) 50%. Spectrum and intensity of light at different heights, *L*, and at different locations inside “light-box” and greenhouse was measured with a fiber-coupled portable Ocean Optics (OCEAN-FX-VIS-NIR) spectrometer.

**Figure 3.**
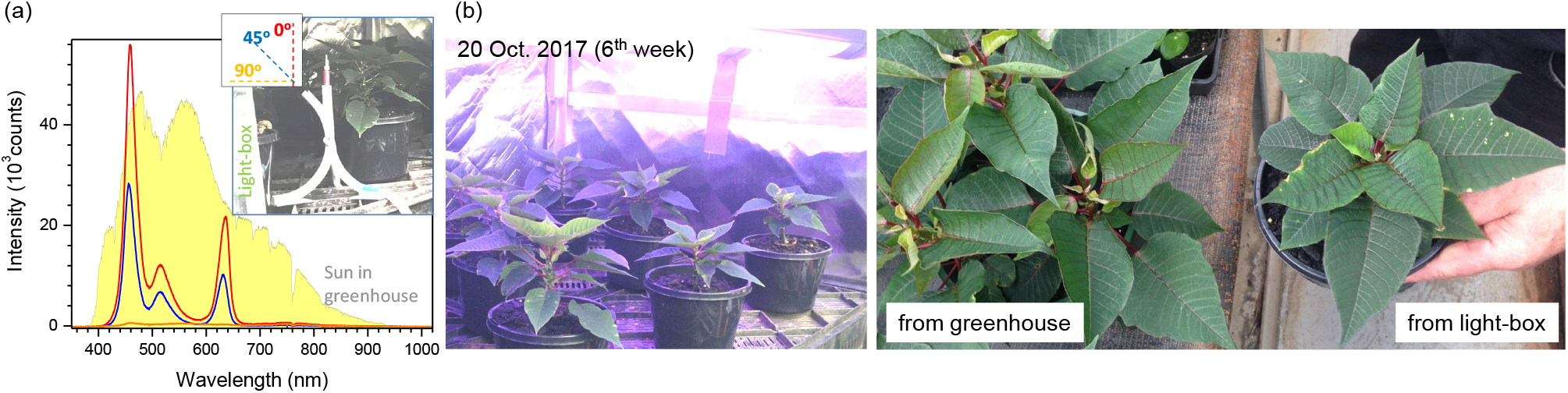
(a) Illuminance spectra in “light-box” at three different orientation angles of the fiber coupler shown in theinset; inset-photo shows 3D-printed stand of fiber-coupler in the measurement position. Illuminance of the Sun in the greenhouse was measured at 0° (up-ward orientation) on 27 Oct. 2017 around 2:00 pm; all spectra taken with integration time of 20 ms. Integrated area difference between Sun and LED spectra is 24.5%; luminance ratio for the LED lamp *I*_45°_/*I*_0°_ = 47.2%. The (b) Poinsettias on 6th week of experiment.

Uniformity of intensity and spectrum over the floor of the “light-box” was measured with separation of ~30 cm (typical distance between plants) at the level of top leaves *L* ≃ 30 cm (Fig. 2). Since the LED lamps with diffusers were used, the uniformity of illumination of the growth area approximately 70 cm from the lamps was uniform within ∼ 15% difference. Interestingly, luxmeter measurements are less informative since it has sensitivity maximised at the green part of the spectrum 500-530 nm where there was much less light emitted from the LED lamp. Luxmeter reading inside the light box was only 1.6% of that in the greenhouse.

Hardware control and monitoring was realised from a personal computer, however, a remote control and data collection can be realised as we have demonstrated in another application of RGBA-lighting for experiments with psychological effects related to color.^8^

### 3.2 Field test: July-December

After 6 weeks of growth (Fig. 3(b)), poinsettias from the “light-box” where slightly smaller. This correlates with an overall light intensity which was smaller for the “light-box”. The greenhouse had illumination by additional eight incandescent 20 W lights for the additional 4 hours per day. Hence, photosynthesis is producing smaller bio-mass. Another contributing factor might be a closer proximity to the light source in the “light-box” which reduces competition for light more expressed in the greenhouse. Figure 4 illustrates this with a side-view image taken when plants entered the final stage of the growth cycle and additional illumination was stopped in the greenhouse on the 21st of November. For the “light-box”, the move from 24/7 illumination to a normal day/night cycle occurred on the 6th of December. A more dense and a better structured plant was formed in the light-box. A side-view also reveals a denser growth while the top-view shows a better structural form for the light-box poinsettia. As a commercial product, the plant grown in the light-box is of a greater value.

**Figure 4.**
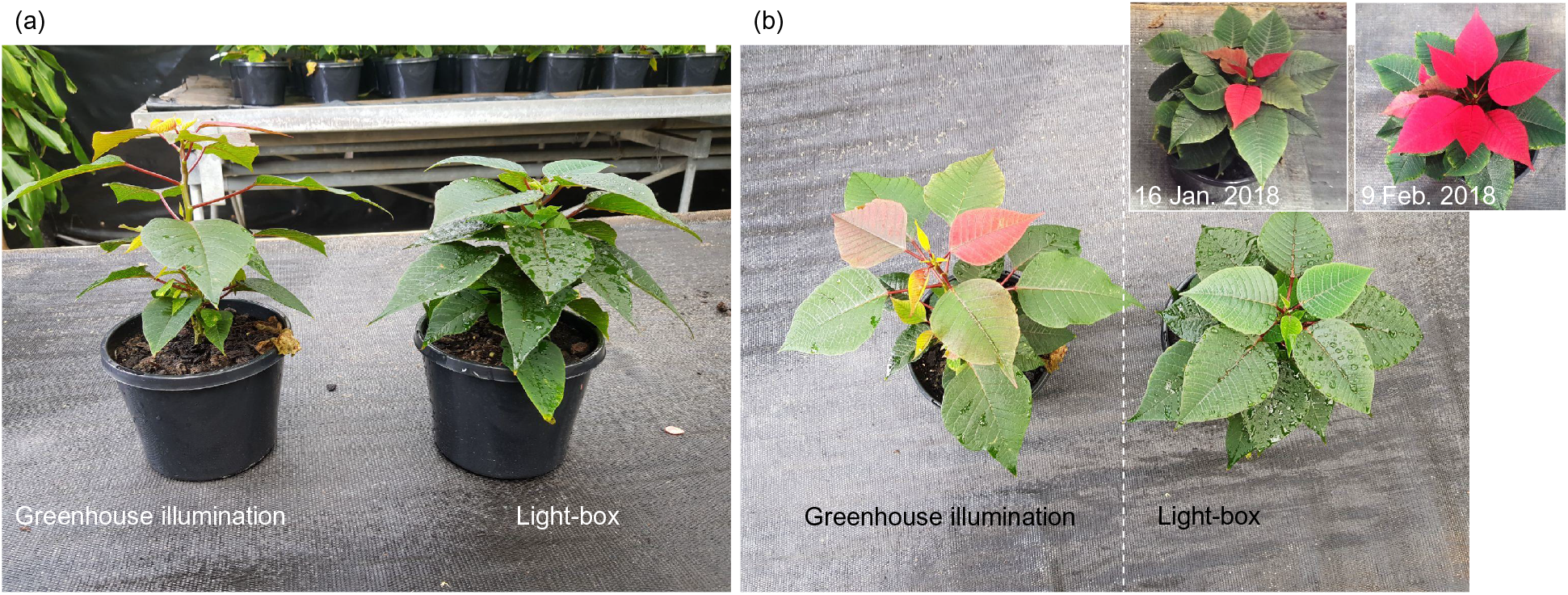
December 6, 2017. (a) Side-view photos reveal that plants were taller with natural greenhouse illumination (which was initially lengthened with halogen lamps for an extra 4 hours and then was switched off for the final stage of leaf coloration). (b) The top-view images for the same plants as in (a). The plant in the light-box had a light totally switched off for the night time, which was applied a two weeks later (on Dec. 6; the day pictures were taken) as compared to the start of a shorter day time illumination in the main greenhouse. The insets in (b) show the plant from the light-box on 16 January and 9 February 2018.

Due to comparatively small enclosure of plants in the “light-box”, plants were slightly more affected by parasites, however, this is a natural consequence of a lower efficiency of ventilation.

## 4. CONCLUSIONS AND OUTLOOK

We have demonstrated a simple do-it-yourself approach in realisation of LED-lamps for agricultural applications. A full cycle of commercial growth of poinsettias during July-December was carried out in the “light-box” conditions. Noteworthy, the illumination cycle transition occurred 3 weeks later for the “light-box” plants; after this transition, the onset of colour change took 1 week longer than their greenhouse counterparts. Luminance in the light-box was less than half that of the greenhouse, however, bio-mass production differed by less than 20%. RGB LED lamp was illuminating plants all the time without day-night cycle for the first 10 weeks.

This work shows that field test using affordable infrastructure is in reach and scientific questions on optimal spectral conditions, circadian cycle control, etc., can be now tested for different decorative and edible plants. The “light-box” could be combined with hydroponics for growth of vegetables in demanding conditions such as encountered in Antarctic research stations where hydroponics are used.

High efficiency and longevity of LEDs with a trend of decreasing price should make this technology commercially viable. By combining artificial LED lighting and quantitative monitoring of photosynthesis with PhotosynQ hand-held technology, new protocols of efficient growth of plants will be developed. It has been already demonstrated that additional illumination by 638 nm wavelength light three days before harvesting of lettuce, marjoram, and green onions resulted in a 44-to-65% reduction of nitrates and increase of nutritionally valuable carbohydrates.^9^

## ACKNOWLEDGMENTS

This work was initiated as part of the technology readiness VG15038 grant: Investigating novel glass technologies and photovoltaics in protected cropping (the work package: LED illumination for the indoor plant growth). This project was supported by Nanotechnology facility of Swinburne University via industrial focus initiative of 2017.

## REFERENCES

[1] Zukauskas, A., Vacekauskas, R., Vitta, P., Tuzikas, A., Petrulis, A., and Shur, M., “Color rendition engine,” Optics Express 20(5), 5356–5367 (2012).

[2] Ledigma Ltd., “Color rendition engine: http://demo.lrg.projektas.vu.lt/lcq/en/simulation/engine, (visited on 22 Oct. 2017).

[3] Morrow, R., “Led lighting in horticulture,” Hortscience 43, 1947–1950 (December 2008).

[4] Choi, H. G., Moon, B. Y., and Kang, N. J., “Effects of led light on the production of strawberry during cultivation in a plastic greenhouse and in a growth chamber,” Scientia Horticulturae 189(Supplement C), 22–31 (2015).

[5] Wojciechowska, R., Dugosz-Grochowska, O., Koton, A., and upnik, M., “Effects of led supplemental lighting on yield and some quality parameters of lamb’s lettuce grown in two winter cycles,” Scientia Horticulturae 187(Supplement C), 80–86 (2015).

[6] Islam, M. A., Kuwar, G., Clarke, J. L., Blystad, D.-R., Gislerd, H. R., Olsen, J. E., and Torre, S., “Artificial light from light emitting diodes (leds) with a high portion of blue light results in shorter poinsettias compared to highpressure sodium (hps) lamps,” Scientia Horticulturae 147(Supplement C), 136–143 (2012).

[7] Embry, J. L. and Nothnagel, E. A., “Leaf senescence of postproduction poinsettias in low-light stress,” Journal of the American Society for Horticultural Science 119(5), 1006–1013 (1994).

[8] Weerasuriya, C., Remote color control of a LED-lamp for human judgement about color (EEE80017 project), Master’s thesis, Nanotechnology facility, Swinburne University of Technology (2016).

[9] Samuolienė, G., Urbonavciūtė, A., Duchovskis, P., Bliznikas, Z., Vitta, P., and Žukauskas, A., “Decrease in nitrate concentration in leafy vegetables under a solid-state illuminator,” Hortiscience 44(7), 1857–1860 (2009).

